# Navigating sex and sex roles: deciphering sex-biased gene expression in a species with sex-role reversal (*Syngnathus typhle*)

**DOI:** 10.1101/2023.05.02.539036

**Authors:** Freya A. Pappert, Arseny Dubin, Guillermo Torres, Olivia Roth

## Abstract

Sexual dimorphism, the divergence in morphological traits between males and females of the same species, is often accompanied by sex-biased gene expression. However, the majority of research has focused on species with conventional sex roles, where females have the highest energy burden with both egg production and parental care, neglecting the diversity of reproductive roles found in nature. We investigated sex-biased gene expression in the broadnosed pipefish (*Syngnathus typhle*), a sex-role reversed species with male pregnancy, allowing us to separate these two female traits. Employing RNA sequencing, we examined gene expression across organs (brain, head kidney, gonads) at various life stages, encompassing differences in age, sex, and reproductive status. While some gene groups were more strongly associated with sex roles, such as stress resistance and immune defence, others were driven by biological sex, such as energy and lipid storage regulation in an organ- and age-specific manner. By investigating how genes regulate and are regulated by changing reproductive roles and resource allocation in a model system with unconventional life-history strategy, we aim to enhance our understanding of the importance of sex and sex role in regulating gene expression patterns, broadening the scope of this discussion to encompass a wide range of organisms.

## 1. Introduction

Males and females of the same species often display sexual dimorphism in morphological traits (1) stemming from sex-biased gene expression (2). Based on which sex displays higher expression levels, genes are categorized as male-biased or female-biased (2). The evolution of sex-biased gene expression can resolve conflicts between males and females, including sexual selection, sexual antagonism, relaxed selective constraint and conflicts in parental investment (1,2). The prevalence of sex-biased gene expression is notable across numerous species, although its variation is assigned to differences across tissues and developmental stages (1–3), known to mostly be lowest during embryonic phases and highest in sexually mature adult stages (4).

Sex-specific gene expression has predominantly been investigated in species characterized by conventional sex roles, encompassing male-male competition and female parental care (1). Given the evolutionary flexibility of sex roles, and their variation among species and across individual’
ss lifetime, research going beyond conventional sex roles is tremendously required for understanding the evolution life-history strategies (5–7). Paternal care is the predominant parental care strategy in teleosts encompassing egg guarding, and oral brooding to full viviparity (7,8). The most devoted male parents can be accredited to the syngnathid family (seahorses, pipefishes and seadragons), which have evolved the unique male pregnancy ranging from simple attachment of eggs to the male ventral side to highly specialized brooding structures, analogous to those found in eutherian mammals, that provide embryos with protection, nutrients, oxygen and immunological components (8–12). Experimental studies on parental investment, however, have focused primarily on the mothers’ fitness, linking it directly to the offspring’s fitness, since females investment into egg production and in caring or nurturing offspring is typically higher (13). The wide diversity of natural sex roles ranging from female-biased parental care with male mating competition to male-biased parental care with strong female competition offers opportunities to explore resource allocation trade-offs as drivers for sex-specific life history strategy (14). Life-history strategy evolved with distinct energy investment into somatic maintenance and reproduction, resulting in differences sex-specific resource allocation for immune defence, metabolism and longevity (15–17). Unfortunately, we are lacking a comprehension of how patterns of sex-biased gene expression fluctuate in species featuring sex-role reversal with parental male-biased care. This basic research is crucial because gender dynamics are changing in human society and evolving over time. Currently, the concept of targeted medicine for sex-specific illnesses primarily relies on the binary classification of biological sex (egg or sperm producer) and tends to overlook the societal and environmental context of that sex. This approach creates a blind spot in biomedical research, as it fails to account for the costs of parental care and gender-specific settings that can result in shifts in gene expression.

To understand the importance of sex vs. sex-role in driving gene expression patterns, we compared sex-biased gene expression of young and older broadnosed pipefish *Syngnathus typhle*. The syngnathid family with their sex-role reversal and their unique male pregnancy evolution facilitates decoupling the role of the female sex (defined as the contribution of eggs) and pregnancy (body reshaping and energy allocation) (11). In the broadnosed pipefish, the females produce on average more eggs than males can brood (18,19), resulting in males being the more reproductively constrained sex (20). Larger brood size and limited resource availability influence the cost of male care, making embryo mortality inversely proportional to male condition (male mortality increases with a larger brood size) (12). Given their polygamous mating system and males being the limited sex, males result in being the more choosy sex whereas females are ornamented and actively court males (11,20,21). In the present study, we sought to investigate how these sex-specific expression patterns vary between young and older pipefish individuals and unravel the influence of age and resource allocation shifts on sex-biased gene expression across distinct organs and conditions. To achieve this, we conducted full transcriptome RNA sequencing (RNA-Seq) to analyse gene expression in three distinct organs: brain, head kidney, and gonads, across different age groups (approximately 1 year old and over 2 years old) and between female and pregnant/non-pregnant male pipefish. Our findings deepen our understanding of this unique system and contribute to discussions about sexual dimorphism, parental investment as drivers of gene expression.

## 2. Methods

### 2.1 Sample collection

Broadnosed pipefish *Syngnathus typhle* were caught in the south-western Baltic Sea (54°39’N; 10°19’E) in spring during their breeding season (22). They were brought to the GEOMAR institute, where body mass and total body length were measured. In our study we used 10 females and 20 males; the latter included 10 pregnant and 10 non-pregnant. We also separated the pipefish based on age, ca. 1 year old for young pipefish and above 2 years for old (their estimated lifespan in the wild is 3 years) and measured body size and weight (Figure S1). Given the limited availability of established models for studying non-model organisms like pipefish, our approach draws on years of fieldwork experience. Broadnosed pipefish engage in mating and breeding activities during spring, followed by migration to unknown areas during winter, before returning to the seagrass meadows of the south-western Baltic Sea for breeding in the subsequent year (22). The offspring cohorts from one breeding season typically only reach sexual maturation during the next breeding season, indicated in males over the development of a brood pouch that only becomes visible during the next breeding season (phenotypic signal for sexual maturation).

The animals were sacrificed with an overdose of MS-222 (Tricaine methane sulfonate, 500 mg/l; Sigma-Aldrich). Carcasses were dissected, and different organs including brain, head kidney, and gonads were sampled and immediately preserved in RNA later. The samples were kept at 4 °C for three days, before being transferred to - 20°C for long-term storage. Previous studies have highlighted substantial variability in sex-biased gene expression among organs (23–25). Gonads, exhibit the highest degree of sex-biased expression, with most biased genes involved in sex differentiation and fertility regulation (1,25). Notably, brain-specific dimorphism has demonstrated preliminary trends in *Syngnathus scovelli* (26), while the head kidney—a unique organ in teleost fish analogous to the adrenal gland—serves the dual function of producing red and white blood cells, thereby supporting both oxygen transport and immune defense (27). Given its prominent role in immune regulation, the head kidney becomes a focal point for studying genes displaying sexual immune dimorphism in expression as identified in diverse species from insects to lizards, birds and mammals, with an overall stronger innate and adaptive immune response in females (28,29).

### 2.2 RNA isolation and Illumina sequencing

For our RNA-Seq analysis, we extracted RNA from brain, head kidney, ovaries and testes. The RNA extraction was performed using RNeasy Mini Kit (Qiagen, Venlo, Netherlands) according to the manufacturer’s protocol. Extraction yields were quantified using NanoDrop ND-1000 spectral photometer (Peqlab, Erlangen, Germany), RNA was then stored at -80°C. Library preparation (TruSeq stranded mRNA Kit by Illumina) and sequencing (RNA-Seq NovaSeq6000 S4, 2x150bp, over 17M clean paired-end reads per sample) were carried out at the Institute of Clinical Molecular Biology (IKMB).

### 2.3 Data analysis

#### 2.3.1 Morphological analysis

Statistical analysis was done in Rstudio (v.4.2.2) (30) and tests were considered significant when p-values were smaller than p= 0.05. Mean and standard deviations were calculated and in a separate two-way ANOVA the differences between weight [g] and length [cm] for sex and age were analysed, with Tukey’s Honestly Significant Difference (TukeyHSD) post-hoc test for single interactions (boxplots for weight and length: Supplementary Material, Figure S1).

### 2.3 Transcriptome analysis

Two samples had to be excluded due to sequencing failure (one YM testes and one OM pregnant testes). The resulting reads were quality-controlled using FastQC v.0.11.9 (31) and trimmed using Fastp v.0.20.1 (32). Reads were aligned to a whole genome assembly of *Syngnathus typhle* (BioProject ID: PRJNA947442) using STAR v.2.7.9a (33). Transcript abundance was quantified with TPMCalculator (34).

Statistical analysis in Rstudio (v.4.2.2) (30) used the edgeR package (v.3.40.2) (35). We scaled the raw count data with 26072 genes to counts per million (cpm) and filtered it based on a minimum threshold of 10 counts in at least 20 libraries, resulting in 19348 genes. To control for composition biases, we normalized the data using the trimmed mean of values (TMM) with the *calcNormFactors* function.

We initially performed a principal Component Analysis (PCA) with the “regularized log transformation procedure” (rld) transformed expression values across all the samples to validate organ separation and check for outliers (Figures 1 and S2). Ellipses on the PCA plots represent 95% confidence intervals (Figure 1B). Once we had assessed that organ type explained most variation in the dataset, we focused on single organ’
ss differential gene expression (DGE) analysis, to detect age, sex and pregnancy effects. The raw count data was separated in brain, head kidney, ovaries and testes. We reapplied filtering and normalisation steps and resulted in 18373 genes in brain, 16746 in head kidney, 16368 in ovaries and 18218 in testes. For each organ we plotted PCAs (Figures S2) and tested our hypotheses using a Permutational Multivariate Analysis of Variance Using Distance Matrices (PERMANOVA) using function “adonis2”, method “bray” and 999 permutations (adonis2 (counts ∼ meta$age*meta$sex*meta$pregnant, permutations = 999, method = “bray”). As a post-hoc we used the *pairwise*.*adonis* function with Bonferroni correction.

**Figure 1:**
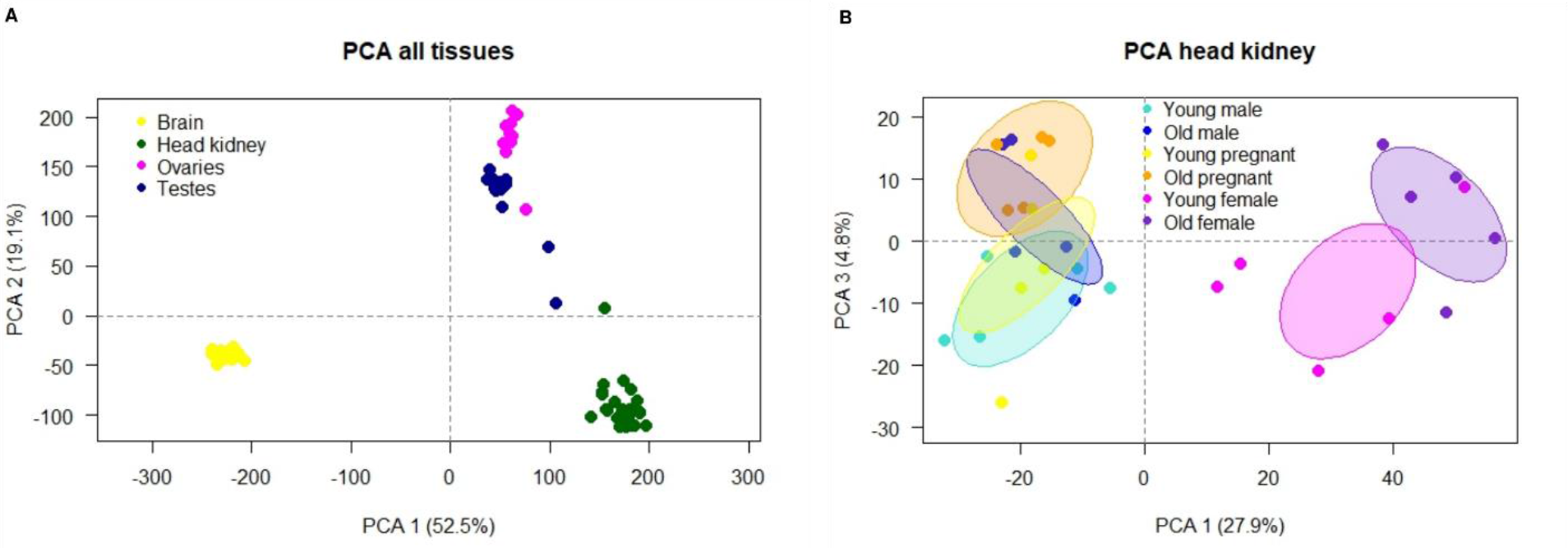
A) PCA plot showing sample clustering for organ type. Each organ is represented by distinct colours: brain in yellow, head kidney in green, ovaries in magenta, and testes in dark blue. B) PCA analysis specifically focusing on head kidney samples. The plot portrays the relationships among different groups within this organ. Six groups are highlighted: young males in light blue, old males in dark blue, young pregnant individuals in yellow, old pregnant individuals in orange, young females in magenta, and old females in purple. The graph emphasizes PC1 and PC3, unravelling significant variations and potential clusters within these groups. Additional information regarding PC2 can be found in the supplementary material (Fig. S2C).

Subsequently, we employed the *limma* package (version 3.54.1) for conducting DGE analysis (36). *limma* employs a linear model approach coupled with empirical Bayes techniques and utilizes the voom method to transform count data into a continuous scale. This method is especially effective in addressing batch effects and reducing technical variability. The voom method is specifically adept at handling varying library sizes (37). To compare multiple groups (Table S2), we constructed a matrix containing independent contrasts. This matrix facilitated conducting a one-way analysis of deviance (ANODEV) for each gene. Following this, we calculated coefficients representing differences between groups to ascertain log fold changes (logFC) through the application of the *contrasts*.*fit* function. Then we implemented empirical Bayes moderation to shrink the estimated variance and conducted moderated t-tests to pinpoint genes with significant differential expression. For subsequent downstream analysis, our focus was exclusively on adjusted P-values below 0.05 and logFC ± 1. This ensured that the observed effects of treatments were statistically meaningful.

#### 2.3.4 Gene ontology and enrichment analysis

For gene ontology analysis, we annotated the differentially expressed genes using a homology-based search of *Danio rerio* (GRCz11) with OrthoFinder (38). *Danio rerio* orthologues from the resulting DEGs between YM vs. YF and OM vs. OF in head kidney were uploaded to g:Profiler web tool (last accessed on 11^th^ August 2023) (39) to perform gene enrichment analysis.

The settings used were: Organism “*Danio rerio* (Zebrafish)”, statistical domain scope set at “All known genes”, significance threshold “g:SCS threshold” at 0.01, numeric IDs treated as “ENTREZGENE_ACC”, the data sources were limited to GO biological processes and biological pathways the databases Kyoto Encyclopedia of Genes and Genomes (KEGG) and Reactome (REAC) (Tables S4). To make our analysis more specific, we applied a maximum term size of 1000 to exclude overly broad categories and prioritize more informative outcomes.

## 3. Results

### 3.1 Sex-biased expression between organs

The results illustrating the variations across distinct organs are depicted in the PCA plot shown in Figure 1A. Notably, the samples exhibited robust clustering based on their organ types, with the brain manifesting the most pronounced distinction compared to the remaining organs. Given that the primary source of variance within the dataset arose from organ types, and our interest was focused on delving into the impacts of sex, age, and pregnancy status, we proceeded to conduct a more granular analysis within each individual organ.

Upon closer examination of the brain, hypothesis testing did not reveal any statistically significant disparities for the studied variables or their interactions (PERMANOVA, P > 0.05). This outcome is also visually evident in the PCA plot specifically dedicated to brain tissue (Fig. S2A). While discernible differences did exist among the sex-specific gonadal tissues (Fig. S2B), a detailed exploration within the testes and ovaries of the two distinct age groups (young vs old) failed to unearth any noteworthy effects. Nonetheless, a modest effect emerged from the interaction between age and pregnancy status (yes or no) within the context of testes (PERMANOVA, P = 0.034). Subsequent pairwise comparisons, however, did not yield any interactions that reached a statistically significant threshold. In terms of head kidney, neither age nor pregnancy displayed substantial impacts, whereas sex exerted a highly significant influence (PERMANOVA, P = 0.001). This distinction is depicted in the PCA plot presented in Figure 1B, wherein PC1 accounts for approximately 28% of the variance between samples. Additionally, PC3 appears to mildly differentiate between young and old individuals, with both OM and OP subjects clustering toward the upper left corner, and OF individuals gravitating in a similar direction on the right, albeit without achieving statistical significance.

The DGE analysis suggested that distinctions pertaining to sex, age, and pregnancy were particularly pronounced within the head kidney (Tables S2 and S3). Notably, in the contrast comparison between YM and YF, 1930 significantly DEGs were identified (adj.p.val < 0.05), while the comparison between OM and OF yielded 3564 DEGs. Nonetheless, no differentially expressed genes (DEGs) were observed when comparing pregnant and non-pregnant males across all organs (Table S2). Results in the brain revealed only a couple of DEGs, which were discarded later in the analysis due to either low logFC or the absence of Danio rerio annotations (Tables S2 and S3).

Conversely, DGE analysis within the respective gonadal organs (testes and ovaries) displayed an absence of significantly DEGs, when considering age or pregnancy (Table S2). Yet, DGE analysis of testes vs. ovaries, revealed a high number of significantly DEGs for age (Tables S2). With a logFC threshold of ±1 we found 6620 DEGs between YM vs YF, with 3370 upregulated in YMs and 3250 in YFs, and 7504 for OM vs OF, with 3574 genes upregulated in OMs and 3930 upregulated in OFs. However, when checking for overlaps of DEGs between the two age groups, we found 5623 DEGs that did not change with increasing size and thus age. Meaning that most genes were sex-biased irrespective of increasing age. Nonetheless, there were roughly 997 significant DEGs between YMs and YFs which were not DE between OM and OF and 1881 vice versa, making these age dependent. GSEA of the *Danio rerio* annotated genes revealed that testes had the most enriched pathways involved in transmembrane transport (monoatomic ion - GO:0034220, inorganic ion - GO:0098660, more in Table S4), translation pathways (GO:0006412, GO:0070588, REAC:R-DRE-72766) and Nonsense-Mediated Decay (NMD) (REAC:R-DRE-927802, REAC:R-DRE-975956, REAC:R-DRE-975957) (Table S4). While in ovaries, we foung pathways enriched for ncRNA metabolic process (GO:0034660) and RNA modification (GO:0009451) (Table S4). On closer inspection of DEGs that were uniquely DE between testes and ovaries in young pipefish, we could not find any enriched pathways (adj.p.val < 0.01). While, for older pipefish gonads we did find Mitogen-activated protein kinase (MAPK) signalling pathway (KEGG:04010) enriched in old testes, compared to DNA repair (GO:0006281) and methylation (GO:0032259) pathways enriched in ovaries of older females (Table S4).

### 3.2. Sex-biased gene expression in head kidney

Results from the DGE analysis in head kidney revealed 912 significant DEGs (with logFC threshold of ±1) in the contrast comparison between YM and YF, and 1556 DEGs between OM and OF (Figure 2A). In the first case, 500 genes exhibited upregulation in YM and 412 in YF, while for the OM vs OF comparison, 780 were upregulated in OM and 776 in OF. Overlaps between sex-biased expressed genes for young and old pipefish found that 291 DEGs were upregulated in both YF and OF, whereas approximately 223 were the same for both YM and OM (Figure 2B). GSEA on the latter, revealed that female-biased genes, regardless of age, displayed enrichment (p < 0.01) in processes such as transport (GO:0006810), peroxisome (KEGG:04146), and the terminal pathway of complement (REAC-R-DRE-166665) (Table S4). In contrast, male-biased genes exhibited enrichment in myeloid cell differentiation (GO:0030099) and hemopoiesis (GO:0030097) (Table S4).

**Figure 2:**
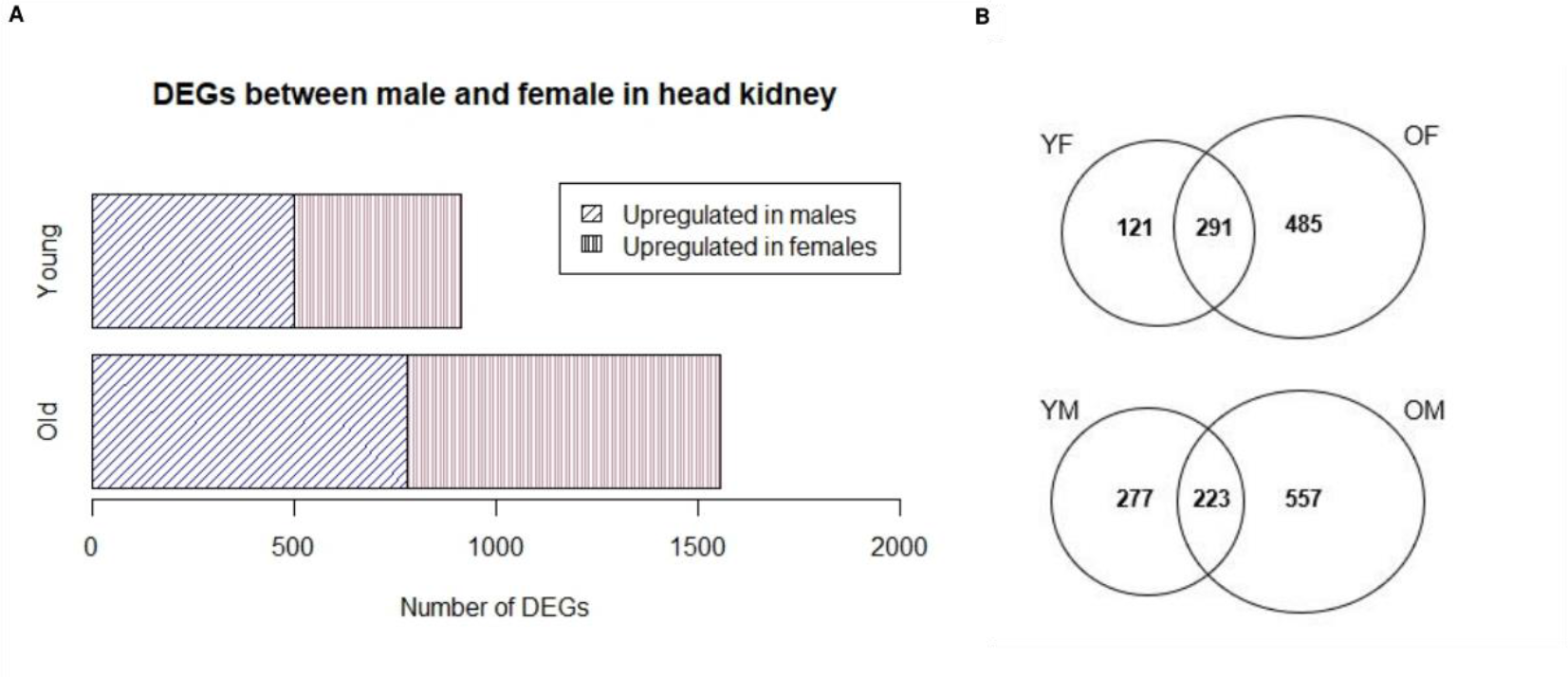
This figure presents the outcomes of the DGE analysis in the head kidney (adj.p.val < 0.05; logFC ± 1), focusing young males (YM) vs. young females (YF), as well as old males (OM) vs. old females (OF). A) Bar plots illustrate the genes that exhibit upregulation in males compared to females, stratified across different age groups. YM display overall 500 upregulated genes vs. 412 in YF, while OM showed 780 upregulated genes vs. 776 in OF. B) Venn diagram showing overlapping sets of DEGs between YF and OF, as well as between YM and OM. This helps us discern genes that exhibit sex-specific biased expression irrespective of age and those whose sex-biased expression changes particularly with age.

Results from the GSEA of significant DEGs that were biased (logFC ±1) in YM or YF, as well as OM or OF, can be visualized in Figure 3 (also detailed in Table S4). Older females exhibited enrichment across categories linked to diverse metabolic pathways, encompassing lipid metabolism (REAC:R-DRE-556833), fatty-acid metabolism (REAC:R-DRE-8978868), and peroxisomal lipid metabolism (REAC:R-DRE-390918). Furthermore, heightened activities in cell respiration and oxidation processes, exemplified by pathways such as peroxisome (KEGG:04146), alpha-oxidation of phytanate (REAC:R-DRE-389599), and biological oxidations (REAC:R-DRE-211859). In contrast, YF predominantly exhibited enrichment in the peroxisome pathway (KEGG:04146) (Table S4).

**Figure 3:**
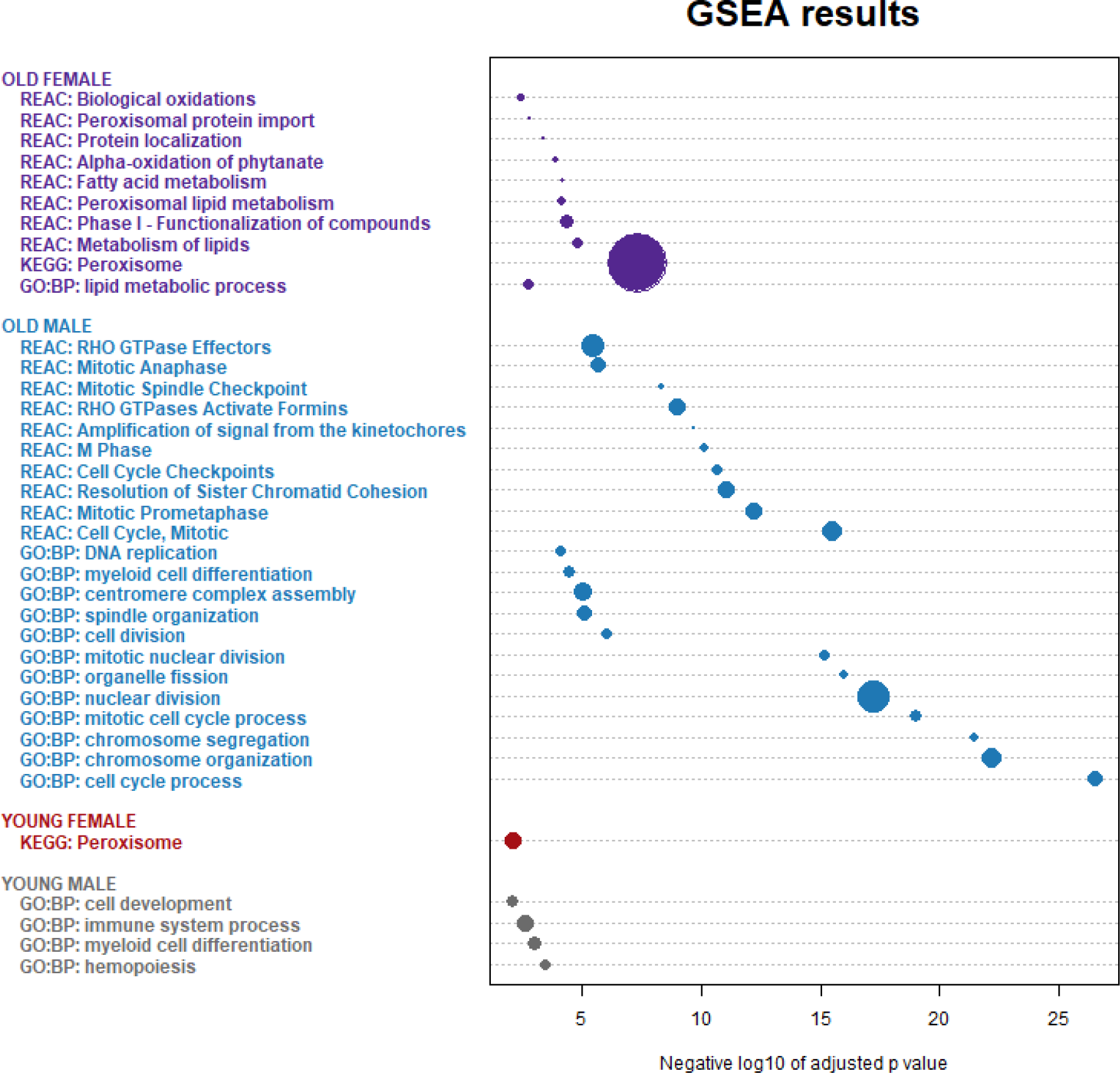
Gene set enrichment analysis results in head kidney, for *Danio rerio* annotated DEGs (logFC < or > 1, adj.p.val < 0.05) upregulated in either YM in gray, YF in red, OM in blue, OF in purple. Data sources were limited to GO biological pathways, KEGG and Reactome, with a g:SCS threshold of 0.01. X-axis shows the negative log10 of the adjusted p values. Sizes of the dots are indicative of the quantitative numerical overall between intersection size and the actual GO term size. As there were numerous pathways highly enriched in OMs, we set a more stringent threshold of adj.p.val < 0.0001 and removed repetitive pathways for easier reading (full results are found in supplementary Table S4).

Genes upregulated in OF compared to OM that were integral to metabolic pathways encompassed several cytochrome P450 enzymes such as *cyp1b1, cyp2r1, cyp4v7, cyp8b1* and *cyp19a1a*, constituting a superfamily of enzymes pivotal in metabolizing a spectrum of compounds, including drugs, toxins, and hormones (40). Their exact function in fish is not known, but for instance, CYP1 localizes to the mitochondrial inner membrane, catalysing the critical initial step in steroid biosynthesis – the conversion of cholesterol to pregnenolone hormones (40). CYP2 enzymes, as recognized in mammals, metabolize vitamin D and arachidonic acid hormones (40). While CYP19, also known as aromatase, facilitates the conversion of C19 androgen to aromatic C18 estrogen (40). Further delving into lipid metabolism, several female-biased expressed genes stood out, including fatty acid amide hydrolase (*faah*), lipoprotein lipase (lpl), and apolipoprotein L 1 (*apol1*), A-II (*apoa2*), and apolipoprotein C-II (*apoc2*). Notably, *faah, apoc2*, and *lpl* also exhibited upregulation in YFs compared to YMs (Table S3). The peroxisome pathway displayed female-biased enrichment in both YF and OF (Figure 3). These enzymes play pivotal roles in fatty acid oxidation and lipid metabolism at large, while also contributing to the detoxification of harmful compounds, including reactive oxygen species (ROS) (41). Particularly pronounced in OF were genes related to cell respiration and oxidation, such as various glutathione peroxidase forms (4a and 4b) and peroxiredoxin (*prdx1*, also observed in YF). Additionally, genes involved in respiration NADPH oxidase 1 (nox1) and 4 (*nox4*), acyl-CoA oxidase 3 (*acox3*, also evident in YF), ferredoxin reductase (*fdxr*), and ferredoxin 1 (*fdx1*, also observed in YF) exhibited upregulation (Table S3).

Numerous pathways exhibited enrichment in OMs relating to cell cycle, chromosome organization, DNA replication, and assorted cellular processes (Figure 3 and Table S4). Some DEGs that overlapped between these pathways belonged to the kinesin family (*kifc1, kif11, kif14, kif15, kif20a, kif20ba, kif22, kif23, kif18a*) or centromere proteins (*cenpn, cenpe, penpk, cenpo, cenph, cenpt, cenpf*, and numerous others) (Table S3). Additionally, still in OM pipefish, pathways concerning DNA repair (GO:0006281, p < 0.002) and DNA damage response (GO:0006974, p < 0.0001) exhibited enrichment (Table S4). Notable genes within these pathways encompassed damage-specific DNA binding protein 2 (*ddb2*) and DNA repair contributors *rad51b* and *rad54*, which play specific roles in homologous recombination-based DNA repair processes (42). Furthermore, ATM serine/threonine kinase (*atm*), checkpoint kinase 1 (*check1*), and checkpoint kinase 2 (*chek2*), with their overarching responsibilities in activating downstream targets implicated in DNA repair and cancer prevention (43,44), were also represented.

Importantly, the timeless circadian clock gene exhibited upregulation in both young and old males compared to females (Table S3). Although fish-specific research in this context is lacking, studies in humans have revealed that its downregulation leads to telomere-associated DNA damage (45,45). Notably, intriguing male-biased expressed genes included *deptor*, a negative regulator of the Mammalian Target of Rapamycin (mTOR) signalling pathway that acts as a feedback inhibitor (46). Additionally, insulin-like growth factor 2 mRNA binding protein 2a (*igf2bp2a*) and, in older males, peroxisome proliferator-activated receptor gamma, coactivator 1 alpha (*ppargc1a*), a homolog of forkhead box O (FOXO) 1, all interconnected in nutrient sensing pathways and collectively influential in cell growth, metabolism, and lifespan regulation (47,48).

For genes particularly upregulated in YMs compared to YFs, we observed enrichment in hemopoiesis (GO:0030097), cell development (GO:0048468), and myeloid cell differentiation (GO:0030099), although these also exhibited enrichment in OMs relative to females albeit to a lesser extent (Table S4). These pathways’ enrichment make sense, given the head kidneys’ role in producing red and white blood cells (27). Interestingly, YMs uniquely exhibited enriched genes associated with immune system processes (GO:0002376) (Table S4). Within this pathway genes included chemokine receptors 1 (*cxcr1*) and 4a (*cxcr4a*), interferon regulatory factor 5 (irf5), and toll-like receptor 8 (tlr8), the latter being a homolog of human tlr1 (49,50). There was also V-set immunoregulatory receptor (*vsir*), analogous to other immune checkpoint molecules, VSIR (VISTA) plays a role in modulating T cell responses, acting as a negative regulator of T cell activation to prevent excessive immune responses and maintain immune homeostasis (51,52). Additionally, the gene cluster of differentiation (CD) antigens, such as *cd7al* and *cd247l*, were identified, which participate in immune recognition processes (53). Notably, recombination activating gene 1 (*rag1*) was also found upregulated in YMs, encoding a protein crucial for V(D)J recombination, this process is fundamental for generating diverse antigen receptor molecules on immune cells, specifically B and T cells (54,55). Moreover, we observed pro-inflammatory cytokines expressed in males, including neutrophil cytosolic factors (*ncf1, ncf2*, and *ncf4*), where NCF1 stands out as a pro-inflammatory protein contributing to the generation of reactive oxygen species that are pivotal for pathogen eradication (56).

Although females did not show enrichment in immune pathways, upon closer observation we found several immune-related genes notably upregulated in both young and old females compared to males. These included interleukins and their receptors, such as il11ra, il17ra1a, il19l, and il10, the latter suppresses the production of pro-inflammatory cytokines such as TNF-alpha (57), and a major histocompatibility complex class I ZCA (*mhc1zca*) (Table S3), which are molecules present on the surface of all nucleated cells and bind to foreign antigens so that CD8+ T-cells can recognize and eliminate the invading pathogens (Garcia et al. 1996). Furthermore, interleukin 6 signal transducer (il6st) and interferon regulatory factor 6 (irf6) were elevated in YFs.

## 4. Discussion

Our work presents a unique exploration into the intricate relationship between male and female life-history traits and their influence on gene expression patterns, particularly in the context of a sex-role reversed species featuring male pregnancy, *Syngnathus typhle*. By conducting comparative analyses of gene expression among males, females, young, old, pregnant and non-pregnant male pipefish across various organs (brain, head kidney, gonads), our aim was to uncover fresh insights into the roles of sex (i.e., egg or sperm producer) and sex-role (i.e., parental investment, mate competition) in shaping sex-biased gene expression patterns.

Our findings highlighted distinct gene expression patterns across different organs, whole brain, head kidney, and gonads. The most abundant DEGs were observed in the gonads and head kidney, with an occurrence of more sex-biased expressed genes between older pipefish, which aligns with research findings in other organisms (4). No significant sex or age differences emerged within brain, which is mostly consistent with observations in other teleost species (23,24). Although one study did find a few sex-biased expressed genes in the pipefish *S. scovelli* (26). Interestingly, pregnancy exhibited minimal or no effect on gene expression across all examined organs. This observation suggests that the physiological changes associated with pregnancy may not significantly affect the selected organs, considering that another study comparing pouch tissues and pregnancy gradients across four different syngnathid species found DEGs in immune pathways and metabolic processes (58,59).

Based on the results and the reproductive biology of *S. typhle*, we suggest that female pipefish may allocate a relatively higher proportion of their resources to pre-copulatory reproduction compared to males. This inference is drawn from the enrichment of female-biased pathways related to RNA modification in ovaries and the observation that older females display enrichment in metabolic pathways associated with lipid and fatty-acid metabolism. These patterns suggest a potential emphasis on oocyte development, maturation, and hormonal regulation, indicating a notable allocation of resources, possibly towards pre-copulatory reproduction (Figure 4). Furthermore, the enrichment of pathways related to DNA repair and methylation patterns, exclusively enriched in ovaries of older females, may indicate an investment in maintaining genomic integrity, which could contribute to ensuring the quality of genetic material passed to the offspring (60).

**Figure 4:**
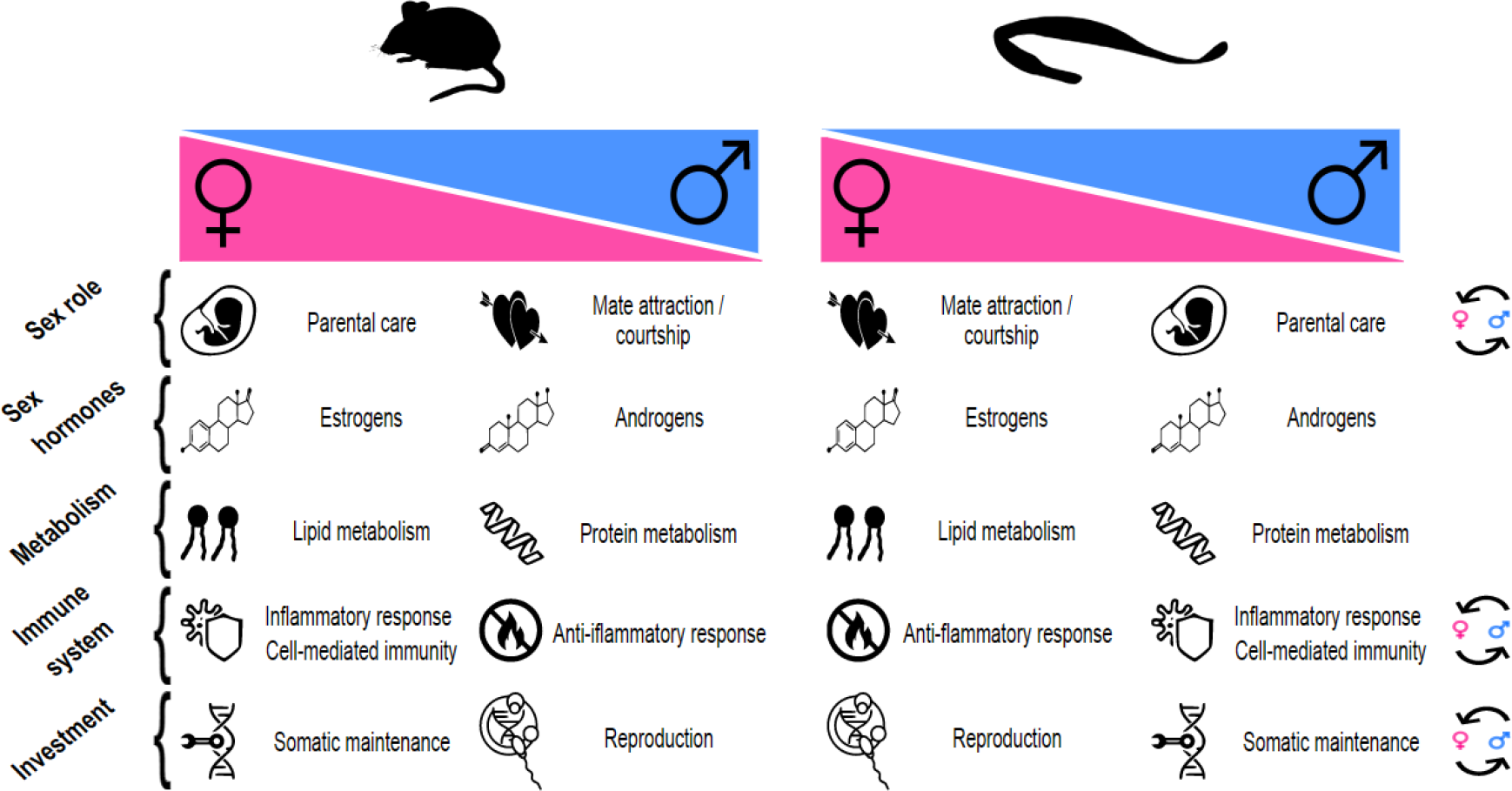
Here represented is a summary comparison of traits between a conventional sex-role mouse model and the broadnosed pipefish, which exhibits sex-role reversal. Specifically, we examine how these traits differ based on biological sex (magenta for females and blue for males). On the far right of the diagram, circular arrows indicate what life-history traits may be reversed compared to the conventional model species.

On a similar note, in the head kidney, female-biased genes, irrespective of age, exhibited a notable enrichment primarily in the peroxisome pathway. These membrane-bound organelles play pivotal roles in lipid metabolism within cells, maintaining cellular oxidative equilibrium (61). Given the energy investment in processes like oocyte development by female pipefish, the heightened peroxisomal pathway enrichment suggests an augmented demand for lipid metabolism to fuel these activities. Notably, older females displayed enrichment in metabolic pathways pertinent to lipid and fatty-acid metabolism (in head kidney), including peroxisomal lipid metabolism. Enriched genes within these pathways, such as *faah* (62), *apolipoproteins* (63), and *lpl* (64), collectively influence lipid metabolism, lipase activity, and energy production. The key role of LPL, as a rate-determining enzyme for fatty acid supply to various tissues, hints at its potential to govern the allocation of dietary lipids towards storage or utilization (64–66).

Regulating dietary fats holds critical importance in fish, serving as both an energy reservoir and a fundamental source of essential fatty acids (EFAs) vital for growth, development, and reproduction (65,67). While basic metabolic processes align between males and females, variations in body composition, sex hormones, sex-specific energy expenditure, and age collectively influence nutrient metabolism and energy storage efficacy (68,69). Notably, oestrogen concentrations influence fat deposition and storage patterns in the body, aligning with findings in *S. typhle* where males produce testosterone and females produce oestrogen despite the reversed reproductive roles (70,71). The involvement of multiple cytochrome P450 enzymes, particularly OFs, such as CYP1 enzyme catalysing the conversion of cholesterol to pregnenolone hormones, which is the starting point in the production of cortisol, oestrogen, progesterone and testosterone (40) and CYP19 helping the conversion of androgen to aromatic oestrogen (oestrone and 17β-oestradiol) (40), implies potential links to oestrogen-driven processes. Thus, the interplay between pre-copulatory energy allocation to eggs and ovarian oestrogen production likely contributes to the sex-biased expression of lipid metabolic-related genes (72).

In contrast, our results suggest that male pipefish, especially older individuals, may exhibit a strategic investment in their health and somatic maintenance. This is supported by the enrichment of the MAPK signalling pathway in the testes, indicating an adaptive response to age-related stressors and changes. The latter is known for its role in various cellular processes like growth, differentiation, and oxidative stress response, this pathway could contribute to sustaining or modulating testicular functions amid aging, possibly by adapting to age-related stressors or changes (73,74). Additionally, in the head kidney of OMs, we found pathways pertaining to DNA repair and damage response, indicating potentially a more robust DNA repair machinery in older males. Male-biased genes such as *deptor, igf2bp2a*, and *ppargc1a*, known for nutrient sensing and pathways related to mTOR, insulin-IGF regulation and FOXO transcription factors, suggests a focus on cellular maintenance, stress resistance, maintenance of cellular homeostasis and potentially, longevity (46–48). The presence of the *timeless circadian clock* gene, associated with genomic integrity and potential telomere-associated maintenance (42), further reinforces the idea of male investment in health. Together, these findings may reflect male pipefish investing in mechanisms that promote their own health and somatic well-being (Figure 4).

An intriguing result was the heightened enrichment of processes related to haematopoiesis and the immune system in YMs as compared to YFs, encompassing genes linked to chemokine receptors, interferon regulatory factors, toll-like receptors, and immune checkpoint molecules. Considering that male pipefish undergo pregnancy and harbour embryos in their pouches, the increased emphasis on haematopoiesis and immune cell development in YMs might signify an evolutionary adaptation aimed at bolstering their own immune system (9). There are inherent trade-offs associated with balancing the reproductive system with the immune system. In species with internal fertilization and gestation, such as pipefish, there are specific demands placed on the immune system, including the need to avoid sexually transmitted diseases and prevent rejection of the developing embryo (75,76). Moreover, the upregulation of *rag1* is noteworthy, as this gene is pivotal for V(D)J recombination—a process generating diverse antigen receptor molecules on immune cells (55). This broader repertoire of immune receptors could effectively recognize a wider array of pathogens. This divergence might reflect evolutionary adaptations that cater to distinct immune system demands between the two sexes (58). We know that sex-specific pathogens exist in other fish species due to sex-specific behaviours, such as shoaling behaviour in female guppies’ increased ectoparasite loads (77). Preliminary in house data suggests that there is a difference in ectoparasite prevalence between male and female pipefish and there is evidence to suggest that parent-specific transgenerational immune priming does occur in broadnosed pipefish (9). The increased number of YM-biased immune genes are in contrast to studies in mammals (Figure 4), where it has been shown that cell-mediated immunity seems to be more pronounced in females with both DC antigen-presenting genes, TLRs and immunoglobulins proliferating at higher rates (29,78). Furthermore, sex-hormone-specific research has shown that androgens typically induce an anti-inflammatory response (29,79), increasing also il10 production, which conversely, was significantly upregulated in female pipefish (having higher oestrogen levels) in our dataset, with also numerous pro-inflammatory markers being upregulated in males (with higher testosterone). The enriched immune pathway in YM might indicate a sex pattern reversal of inflammatory response compared with conventional sex-role species (Figure 4), thus making pregnancy and the life-history trade-offs play a more significant role here. It was surprising however, that we did not find the same immune system pathway enriched in OM and that there were fewer sex-biased immune genes between OM and OF. Aging is associated with a decline in immune function, often referred to as immune senescence. This decline tends to affect both males and females similarly and can lead to a more uniform immune response among older individuals (80,81). This could in part explain our results, but further research is needed.

Our results illuminate that sex-biased gene expression is a dynamic interplay influenced by both biological sex and sex-specific life history traits. Within this intricate framework, resource allocation trade-offs into parental care can exert profound impacts on the patterns of sex-biased gene expression. Notably, our study suggests that some genes may be more influenced by biological sex, while others are shaped by sex roles. While these findings provide insights into potential differences in resource allocation between the sexes, it is important to note that direct quantification of reproductive investment requires additional research, such as assessments of reproductive effort or energy allocation. Furthermore, our research reveals that the gene expression landscape undergoes significant changes with age. Males and females exhibit distinct gene expression profiles, with some pathways converging (e.g., in the pipefish immune system) and others diverging (e.g., in anti-aging mechanisms) over time. These age-related shifts underscore the importance of studying sex-biased expression also in context of senescence and its distinct physiological changes between males and females. The interplay between biological sex, sex roles, age, and organ-specific factors can yield diverse and context-dependent gene expression patterns. In future research, we aim to extend our research beyond a single species of syngnathid, given the diverse array of pregnancy gradients, sex roles, and mating systems present within the family (21,82). We hope to have underscored the importance for a nuanced approach to the study of sex-specific gene regulation, accounting for the multifaceted nature of sex and sex roles in shaping gene expression dynamics.

## Ethics statement

Work was carried out in accordance with German animal welfare law and with the ethical approval given by the Ministerium für Energiewende, Landwirtschaft, Umwelt, Natur und Ditgitalisierung (MELUND) Schleswig-Holstein (permit no. V24257982/2018). No wild endangered species were used in this investigation.

## Authors’ contributions

F.A.P. and O.R. initiated and designed the study. F.A.P carried out the experiment, analysed the data, interpreted the results and wrote the article. G.T. helped develop the DEGs *limma:voom* pipeline analysis. A.D. performed the orthology analysis and provided help with the GO profiling. O.R. helped during the field work, supervised the project and supported the writing of the manuscript.

## Acknowledgements

Special thanks to Dr. Ralf Schneider, who provided useful insights and feedback on the statistical analysis and differential gene expression analysis.

## Data Availability Statement

Supplemental information include additional figures (Figure S1-S2), morphology measurements (Table S2), the significant differentially expressed genes (Table S2-S3) and the gene set enrichment analysis results as tables (Table S4). The raw sequencing data and metadata used in this study is available from the National Center for Biotechnology Information (NCBI) Sequence Read Archive (SRA) under BioProject ID PRJNA943164 with submission ID SUB12940929. The whole genome assembly of *Syngnathus typhle* can be found under BioProject ID PRJNA947442 with submission ID SUB12974183.

