## Supplementary material for "Navigating sex and sex roles: deciphering sex-biased gene expression in a species with sex-role reversal (*Syngnathus typhle*)": Supplemetary files (Figures S1, S2, and Table S1)

### Supplementary Figure S1

Morphological measurement comparison between males (blue) and females (pink) of wild-caught *Syngnathus typhle*. (A) Weight in grams on y-axis and age group on x-axis. (B) Total body length in centimeters on y-axis and age groups on x-axis.

There were significant size differences between young and old. The average weight of the sampled pipefish was 0.65 g (SD = 0.146) in young females (YF) and 0.6 g (SD = 0.082) in young males (YM), whilst 1.71 g (SD = 0.191) in old females (OF) and 1.63 g (SD = 0.243) in old males (OM.). The average total body length was 12.3 cm (SD = 1.02) for YF and 11.56 cm (SD = 0.682) for YM., 16.66 cm (SD = 0.764) in OF and 16.82 cm (SD = 1.41) in OM. A two-way ANOVA of body weight and also of length, for sex and age, showed a significant difference for age (P < 0.0001), but not for sex or the interaction.

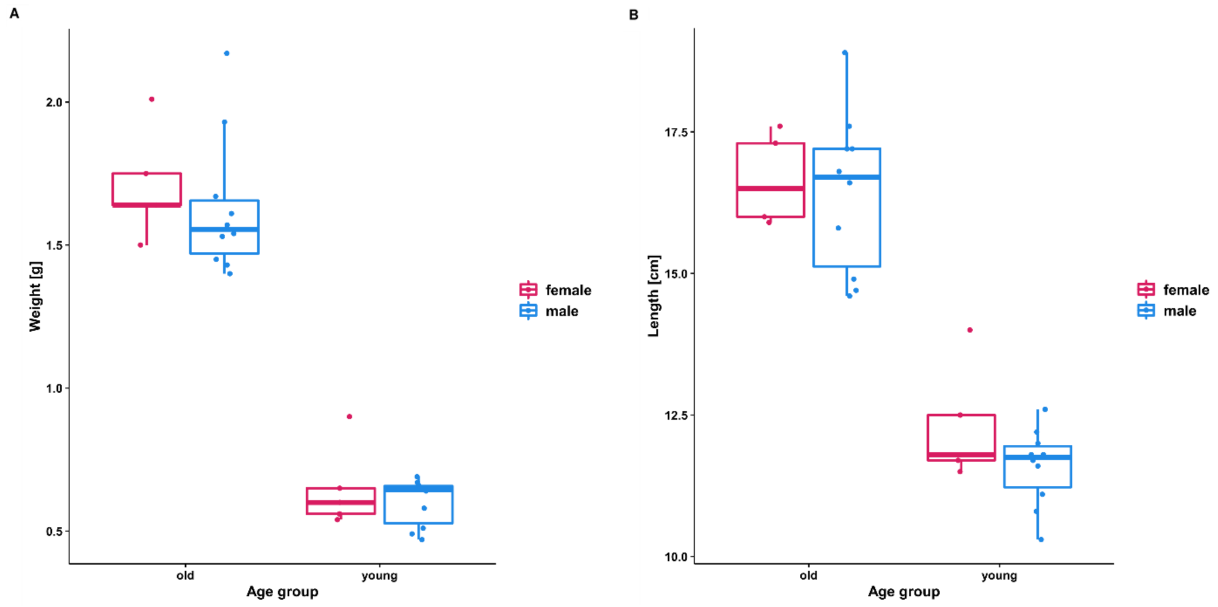

### Supplementary Table S2

Quantitative results from the differential gene expression (DGE) analysis (using *limma:voom*) using contrast comparisons between the different variables for the various organs. Only differentially expressed genes with an adjusted P value smaller than 0.05 were considered (one-way analysis of deviance).

| Contrast comparison | Tissue | Counts (adj.p.val < 0.05) |
|---|---|---|
| YM vs OM | Brain | 0 |
| YF vs OF | Brain | 0 |
| YP vs OP | Brain | 0 |
| YM vs YF | Brain | 2 |

| | | |
|---|---|---|
| OM vs OF | Brain | 2 |
| YM vs YP | Brain | 0 |
| OM vs OP | Brain | 0 |
| YM vs OM | Head kidney | 0 |
| YF vs OF | Head kidney | 4 |
| YP vs OP | Head kidney | 0 |
| YM vs YF | Head kidney | 1930 |
| OM vs OF | Head kidney | 3564 |
| YM vs YP | Head kidney | 0 |
| OM vs OP | Head kidney | 0 |
| YM vs OM | Testes | 0 |
| YP vs OP | Testes | 0 |
| YM vs YP | Testes | 0 |
| OM vs OP | Testes | 0 |
| YF vs OF | Ovaries | 0 |
| YM vs YF | Gonads | 9166 |
| OM vs OF | Gonads | 9493 |

### Supplementary Figure S2

Principal Component Analysis (PCA) results of PC1 and PC2 for the different organs: A) brain; B) ovaries and testes; C) head kidney. The plots portray the relationships among different groups within the respective organ. Six groups are highlighted: young males in light blue, old males in dark blue, young pregnant individuals in yellow, old pregnant individuals in orange, young females in magenta, and old females in purple.

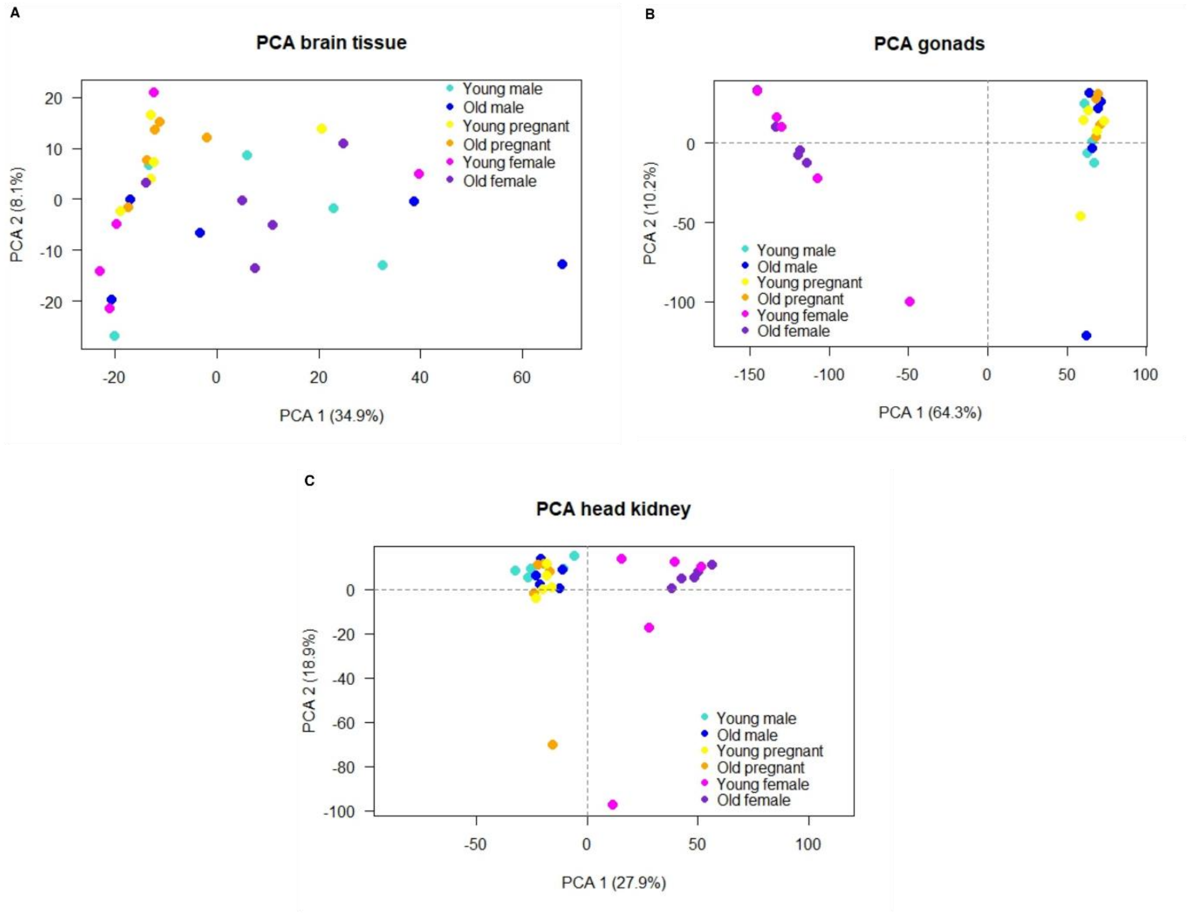
